# Inner Hair Cell and Neuron Degeneration Contribute to Hearing Loss in a Dfna2-Like Mouse Model

**DOI:** 10.1101/469676

**Authors:** Camila Carignano, Esteban Pablo Barila, Ezequiel Ignacio Rías, Leonardo Dionisio, Eugenio Aztiria, Guillermo Spitzmaul

## Abstract

**HIGHLIGHTS:**

- KCNQ4 knock-out mouse shows hair cells and spiral ganglion neuron degeneration.
- Inner hair cells and spiral ganglion neuron loss begin 30 weeks later than outer hair cells in *Kcnq4*^-/-^ mice.
- Inner hair cell loss kinetic is faster than that of outer hair cells in cochlear basal turn in *Kcnq4*^-/-^.
- Outer hair cells from *Kcnq4*^-/-^ mice degenerate slower in apical than in basal turn.
- *Kcnq4* knock-out allele expressed in C3H/HeJ strain reproduces the two phases of DFNA2 hearing loss.

**GRAPHICAL ABSTRACT:** 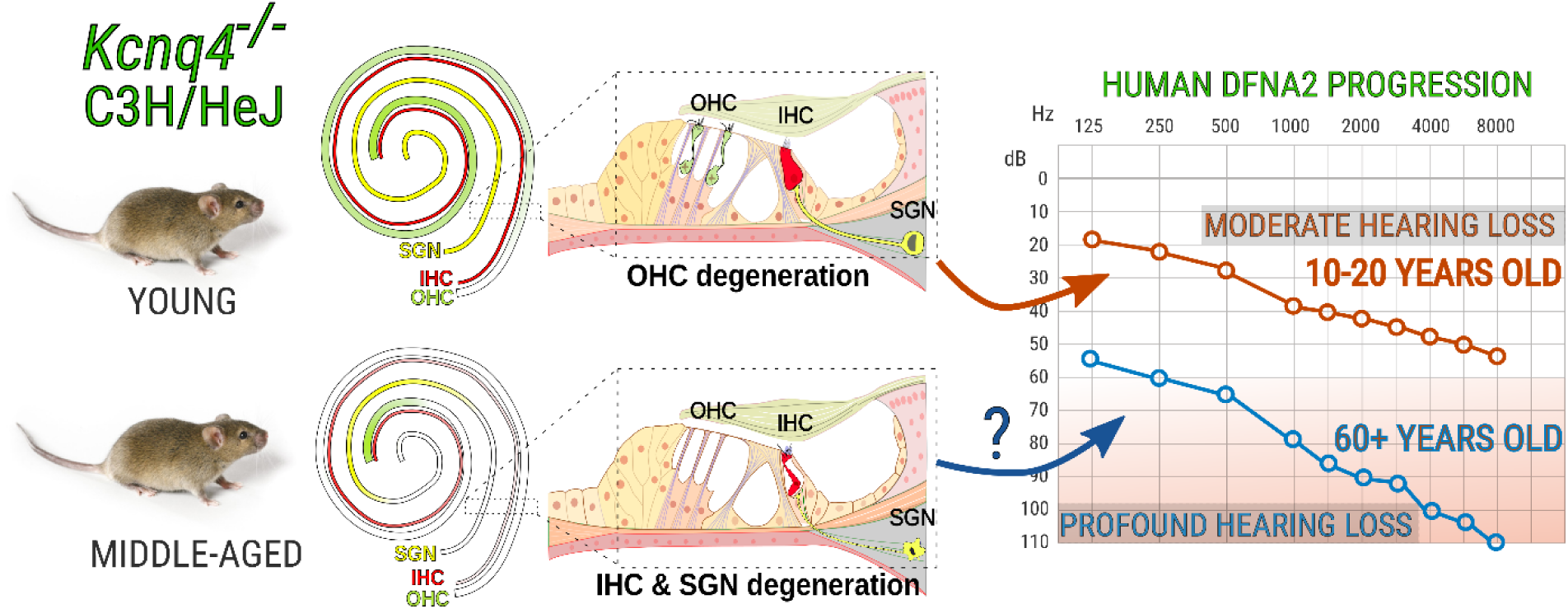

## INTRODUCTION

Potassium circulation is essential for signal transduction in the hearing process (Hibino and Kurachi, 2006, Zdebik et al., 2009), which starts with opening of mechanosensitive cation channels placed at the tips of hair cells (Kurima et al., 2015, Pan et al., 2018). Specialized epithelia on the stria vascularis produce potassium-enriched endolymph that fills in the scala media. To maintain cell homeostasis, once potassium enters hair cells (HCs), it must return to the stria vascularis by exiting HCs and then moving through supporting cells of the organ of Corti (OC). This potassium circulation ensures that the physiological processes of hearing can take place (Hibino and Kurachi, 2006, Mistrik and Ashmore, 2009). Alterations in any of the steps involved in this circulation process can generate different kinds of hearing loss (Van Laer et al., 2006, Lang et al., 2007). KCNQ4 is the main K^+^ channel involved in potassium extrusion from outer hair cells (OHCs). It is a voltage-activated channel localized at the basal pole of OHCs, generating the I_K,n_ current (Kharkovets et al., 2000, Kharkovets et al., 2006). Its function is essential for cell survival. Mutations in KCNQ4 cause disruptions on potassium recycling and its consequences are displayed in the DFNA2 hearing loss (HL) (Kharkovets et al., 2006). This last is a slow progressive deafness characterized by having two phases. The first one starts at around 15-20 years-old exhibiting a mild to moderate HL at high-frequency sounds that progresses over time to middle frequencies (Dominguez and Dodson, 2012). Hearing threshold increases 20 to 60 dB which correlates with a progressive loss of the amplificatory function exerted by OHCs (Nie, 2008). The second phase is characterized by the progression of HL to a severe impairment by the age of 65-70 (De Leenheer et al., 2002). At this stage, hearing threshold increases further beyond 70-80 dB affecting most frequencies which cannot be explained by OHC dysfunction alone (Dominguez and Dodson, 2012). Therefore, other mechanisms must be contributing to the pathology progression. Patients with this condition are heterozygous for the affected *Kcnq4* allele which bears a point mutation with dominant-negative effect on the tetrameric channel (Kubisch et al., 1999, Dominguez and Dodson, 2012). KCNQ4 channel is not only located in OHCs, but also in inner hair cells (IHCs) and some neurons from central nervous system (CNS) nuclei of the brainstem belonging most of them to the auditory pathway (Beisel et al., 2000, Kharkovets et al., 2000, Oliver et al., 2003, Beisel et al., 2005). For these reasons, it is believed that IHCs and neurons could also participate in the progression of HL (Dominguez and Dodson, 2012). Many years have passed since the molecular cause of DFNA2 was discovered (Kubisch et al., 1999). However, neither the molecular events gated by KCNQ4 misfunction are fully understood nor the role of IHCs and neurons on disease progression. Research on DFNA2 echoes into the physiology of normal hearing and the comprehension of several hearing pathologies like age-related hearing loss (ARHL) and noise-induced hearing loss (NIHL) that share alterations in potassium circulation (Fransen et al., 2003, Van Laer et al., 2006, Dominguez and Dodson, 2012, Wong and Ryan, 2015).

Then, in order to understand the contribution of KCNQ4 to potassium homeostasis in the OC we used a KCNQ4 knock-out (KO) mouse that resembles many of the characteristics observed in DFNA2 disease. Remarkably we found that in the absence of KCNQ4, not only OHCs are affected but also IHCs and SGNs. Time-course of cell death is differentially developed for each cell type, suggesting distinct roles of KCNQ4 in these cell types. Our results shed light on the mechanisms that would participate in the progression of DFNA2 deafness.

## EXPERIMENTAL PROCEDURES

### Animals

C3H/HeJ transgenic mice, lacking the expression of the KCNQ4 protein (*Kcnq4*^-/-^), due to a deletion spanning exon 6 to exon 8, were used (Kharkovets et al., 2006, Spitzmaul et al., 2013). Wild-type (WT) C3H/HeJ litters and C57BL/6 mice were used as controls (*Kcnq4*^*+/+*^) and for inter-strain comparison, respectively. Mice from both sexes were employed in all experiments indistinctly. Age ranges were: a) young mice: 3, 4, 6, 8 and 10 postnatal weeks-old (W); b) middle age adult mice: 40W, 52W and 58W. The experimental protocol followed in this study was approved by the Council for Care and Use of Experimental Animals (CICUAE, protocol N° 083/2016) of the Universidad Nacional del Sur (UNS), whose requirements are strictly based on the European Parliament and Council of the European Union directives (2010/63/EU).

### Tissue preparation

Mice ranging from 3W to 58W were euthanized by CO_2_ exposure and inner ears were promptly removed from temporal bones. In order to monitor hair cell and spiral ganglion neuron degeneration, cochleae were studied by immunofluorescence using two different approaches i) mounted as a whole or, ii) in thin tissue sections. Both started with tissue fixation by overnight submersion in 4% paraformaldehyde, washed with PBS, and decalcified using 8-10% EDTA in PBS for up to 5 days depending on animal age, on a rocking shaker at 4°C.

### Whole-mount cochlear preparations

The organ of Corti from decalcified cochleae was obtained following a protocol similar to that described in Akil and Lustig, 2013 and Montgomery and Cox, 2016 (Akil and Lustig, 2013, Montgomery and Cox, 2016). This method consists in splitting the whole cochlear length into three longitudinal segments: basal, middle and apical turns using fine scissors. Then, the vestibular system, spiral ligament, modiolus and tectorial membrane were removed and the organ of Corti was isolated. Finally, the hook was cut off from the rest of the basal segment.

### Modiolar sections

Whole inner ears were processed according to Spitzmaul et al, 2013 and Barclays et al. 2016 (Barclay et al., 2011, Spitzmaul et al., 2013). Briefly, after decalcification, inner ears were cryoprotected in 15% sucrose for 4 h, then in 30% sucrose overnight followed by OCT embedding. 10 μm-thick sections, longitudinal to the modiolus, were obtained using a cryostat (Leica CM 1860) and preserved at −20°C until processed.

### Immunofluorescence in whole-mount cochlear preparations

Cochlear turns were postfixed in 4% PFA during 30 min, washed three times in PBS and incubated for 2 h in blocking solution (2% BSA, 0.5% Nonidet P-40 in PBS). Primary antibodies were incubated for 48 h in carrier solution (PBS containing 1% BSA, and 0.25% Nonidet P-40). Subsequently, tissue was rinsed three times in PBS. Secondary antibodies, diluted in carrier solution, were incubated for 2 h at room temperature. After that, samples were washed three times in PBS. Finally, cochlear turns were mounted unflattened in Fluoromount-G (Southern Biotech). The following primary antibodies were used: rabbit anti-KCNQ4 (K4C, 1:200; generously provided by Dr. T. Jentsch, Forschungsinstitut für Molekulare Pharmakologie, Berlin, Germany), goat anti-prestin (1:200, cat#sc-22692, Santa Cruz Biotechnology), rabbit anti-myosin VIIa (1:200, cat#25-6790, Proteus Biosciences). The following fluorescently-labeled secondary antibodies were obtained from Molecular Probes and used diluted (1:500): donkey anti-rabbit 555 (cat#A-31572) and donkey anti-goat 488 (cat#A-11055). Nuclei were stained with DAPI (1:1000).

### Hair cell counting and cytocochleogram plotting

Labeled cells on whole-mount cochleae were imaged using an epifluorescence microscope (Nikon Eclipse E-600) coupled to a CCD camera (Nikon K2E Apogee) and a laser spectral confocal microscope (Leica TCS SP2). Pictures were analyzed using the Image J software and cell counting was carried out manually. HCs were identified based on their tissue localization in the OC. For OHC, row 1 was considered as the innermost cell line followed by row 2 corresponding to the middle line of cells and row 3 to the outermost cell line. Cellular degeneration was evaluated on a cytocochleogram (4-5 animals for each age and genotype). A cytocochleogram is basically a cartographical approach used for determining HC number and distribution along the entire cochlear length (Viberg and Canlon, 2004, Muller et al., 2005, Boyce et al., 2010, Sanz et al., 2015). It is constructed by plotting the number of OHCs or IHCs versus the relative distance to the apex. In order to normalize the OC´s length, cells were counted separately in 20 segments (S), each of them spanning 5% of the total cochlear length, which was considered the whole (100%) distance. Each S was numbered according to the spanning percentage of the total length, starting from the apex (e.g. from 0% to 5%=S5), to the basal end (S100). Apical, middle and basal turns were defined as those spanning S5 to S35, S40 to S70 and S75 to S100, respectively.

### Cell loss rate estimation

The dynamic of cell loss was calculated plotting the average number of living cells in each segment vs. mice age on a logarithmic scale. Data were fitted through the SigmaPlot 12.0 software (Systat Software Inc.) using the Hill equation as follows:

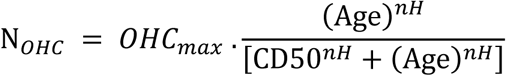

where N_OHC_ represents the number of cells at each age (in weeks), OHC_max_ is the initial OHC number, CD_50_ is the age where 50% cell death was achieved, and nH is the Hill coefficient that estimates cell death rate.

### Immunofluorescence on cochlear sections

Slide-mounted tissue sections were post fixed with 2% PFA for 20 minutes, washed with PBS and blocked in the same buffer as described in section 2.3. Slice sections were incubated overnight with a primary antibody rabbit anti-beta III Tubulin (tuj1, Covance #mrb-435p) diluted 1:1000 in carrier solution. A fluorescently-labeled goat anti-rabbit 488 (Molecular Probes, cat#A-11034) was used as secondary antibody, which was incubated during 1 h.

### Spiral ganglion neuron density measurements

10 μm-thick cochlear longitudinal slices, comprising the whole extension of the modiolus, were chosen to perform SGN assessment. Using a 20x magnification objective, fluorescent images were taken for each of the ganglion individual portions (i.e.: basal, medial and apical). Images were processed with Image J’s FIJI by demarcating their area and taking into account the bound space of bone surrounding the ganglia and the extent of beta III tubulin staining. Using the cell counter extension of this software, individual cells were counted manually and their density was calculated by dividing the number of SGNs over the area in square micrometers covered by the ganglion. Results were plotted for each portion of the ganglion versus the age, in weeks, for each mouse genotype.

### KCNQ4 signal intensity

The expression pattern of the KCNQ4 channel in OHCs was analyzed on whole-mount cochlear preparations from WT mice at different ages. To do so, the organ of Corti was divided into 3 segments, as explained above. To quantify the differences, we measured the mean fluorescence intensity (MFI) of each cell in confocal images using Image J software. For each turn, the intensity of around 20 cells from all 3 rows of OHC in the central area was measured. Background noise was calculated from averaging the signal of 5-6 representative areas where no immunofluorescence was apparent. Specific signal was calculated by subtracting the averaged background value from that of each immunopositive cell.

### RNA extraction and qPCR

Cochlear RNA was extracted from 3-week-old mice. For each experiment, samples from 3-4 mice were pooled. In brief, immediately after cochlear excision, tissue was immersed in a RNA preservation buffer and then, the vestibular region was removed to keep only the cochlea. Using a scalpel, the cochlea was cut between the first and second turn, in order to obtain a bottom piece containing the basal turn and another one containing the middle and apical turns. RNA was extracted using the TransZol reagent (TransGen Biotech, ET101-01) in combination with Direct-Zol RNA mini prep kit (Zymo Research, R2052), essentially as indicated in Patil et al (Vikhe Patil et al., 2015). cDNA was produced from 500 ng total RNA with EasyScript Reverse Transcriptase (TransGen Biotech Cat #AE101) using anchored oligo (dT)s following manufacturer’s indications. Quantitative PCR (qPCR) was carried out using the cDNA generated previously employing the SensiFAST SYBR mix No-Rox Kit (Bioline) in a Rotor-Gene 6000 real-time PCR cycler (Corbett Research). Primers for the *KCNQ4* gene were: 5’-TATGGTGACAAGACGCCACAT-3’ (forward) and 5’-GGCAGAAGCACTTTGAGAAGC-3´ (reverse) while for the reference genes were as follows: HPRT 5’-GTTCTTTGCTGACCTGCTGGA-3’ (fwd) and 5’-AATGATCAGTCAACGGGGGA-3’ (rev); GAPDH 5’-GAGAAACCTGCCAAGTATGATGAC-3’ (fwd) and 5’-CCCACTCTTCCACCTTCGAT-3’ (rev). Data analysis was done applying the ΔΔCt method (Livak and Schmittgen, 2001) to obtain relative mRNA expression.

### Scanning electron microscopy (SEM)

Basal cochlear segments were dissected from *Kcnq4*^+/+^ and *Kcnq4*^-/-^ at 4W and 6W. Cochleae were fixed using 2.5% glutaraldehyde for 2 hours and then, decalcified as mentioned previously. Samples were dehydrated with ethanol and a critical point drier was applied. Finally, tissues were imaged with a LEO-EVO 40 XVP-EDS Oxford X-Max 50 scanning electron microscope provided by the Microscopy Facility from Centro Científico Tecnológico – Bahía Blanca (CCT-BB) / CONICET.

### Statistical Analysis

All data and results were confirmed through at least, three independent experiments. Shown values represent the means ± SEM. Significant differences were identified using, i) Student’s t-test for experiments shown in Fig. 1A, Fig. 2, Fig. 3B and Fig. 7 and ii) one-way ANOVA, post hoc Tukey Multiple Comparison test for experiments shown in Fig. 1C, Fig. 4 and Fig. 6. Statistical analyses were performed using Excel and SigmaPlot 12.0 Software. Statistical significance is represented with (*) when p<0.050; (**) when p<0.010 and (***) when p<0.001.

**Figure 1.**
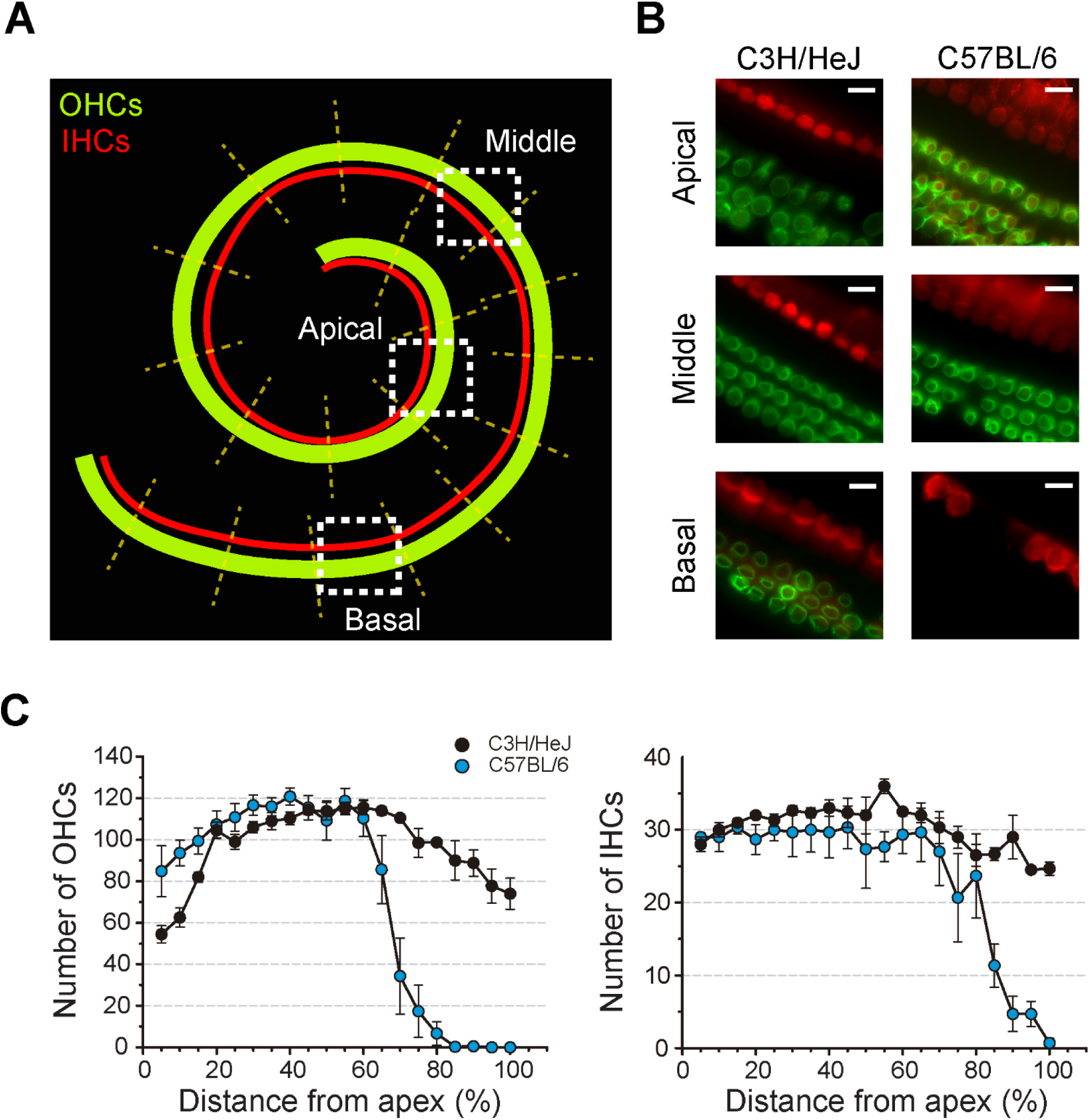
Hair cell survival comparison between C3H/HeJ and C57BL/6 middle-age adult mice. **(A)** *Left*: representative pictures showing hair cell presence in apical, middle and basal turns in C57BL/6 and C3H/HeJ mice at 52W. IHCs and OHCs were labeled with anti-prestin (green) and anti-myosin VIIa (red) antibodies, respectively. Scale bar: 10 µm. *Right*: schematic representation of a mouse Organ of Corti showing its whole extension. OHC (green) and IHC (red) cochlear localization are depicted. Dashed white squares delimit the areas represented on the left panel. Dashed yellow lines indicate segments spanning 5% of the entire cochlear length (S) that were used to build cytocochleograms. Each S was referred relative to the apex. **(B)** Comparison of hair cell degeneration between C3H/HeJ and C57BL/6 mice at 52W. Cytocochleograms showing OHC (left) and IHC (right) counts for each S in C3H/HeJ (black) and C57BL/6 (light blue) strains. Data are represented as means ± SEM (n=3).

**Figure 2.**
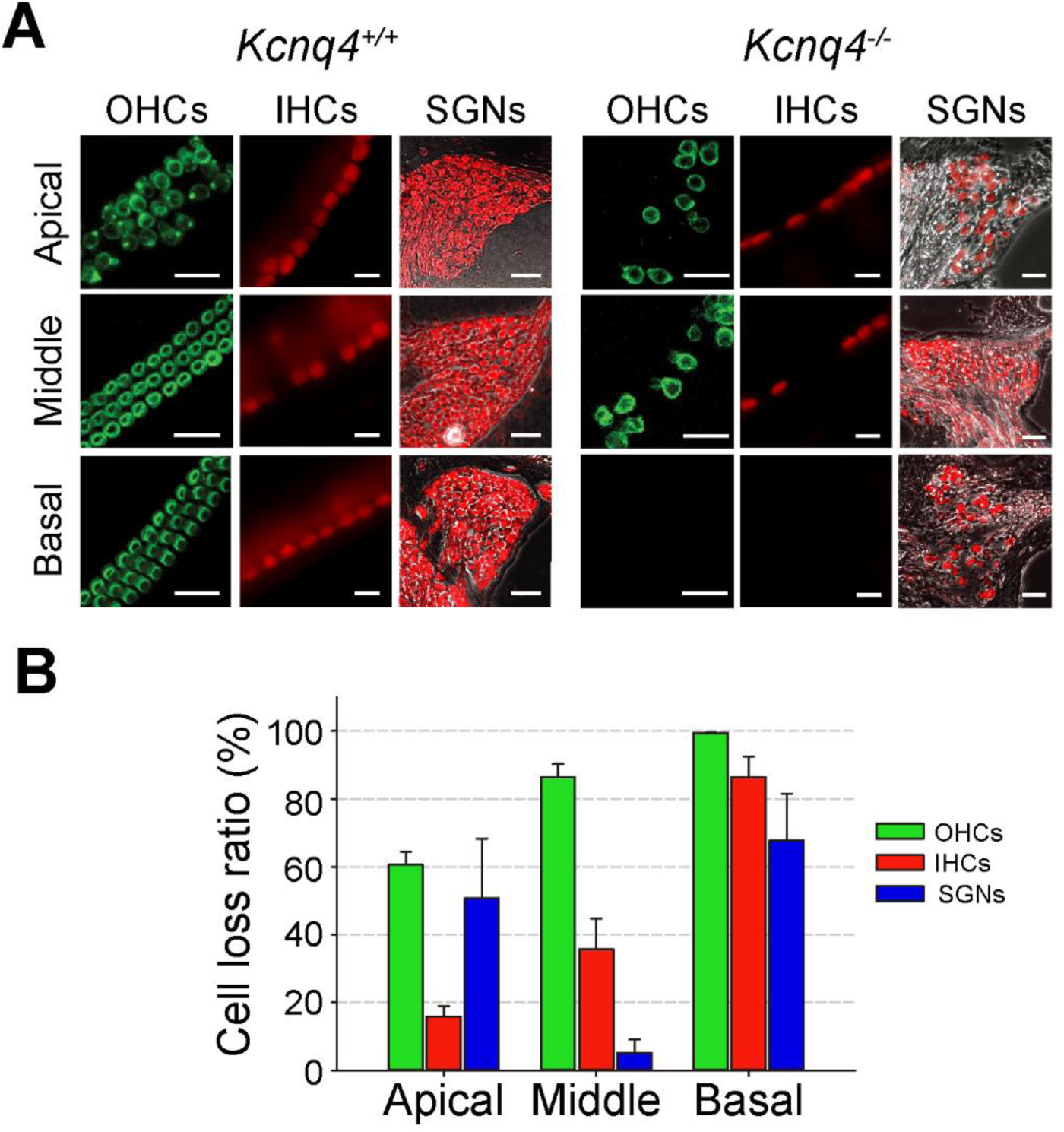
Differential impact of KCNQ4 channel absence on cochlear cell survival. **(A)** Pictures from apical, middle and basal turns showing OHCs, IHCs and SGNs in *Kcnq4*^*+/+*^ and *Kcnq4*^*-/-*^ mice at 58W. OHCs and IHCs were labeled as described in Fig. 1. SGNs were identified with anti-β-III tubulin (red). Scale bars for OHCs, 20 µm; IHCs, 10 µm and SGNs, 50 µm. **(B)** Bar plot depicting normalized cell loss. Cell death ratios for OHCs (green), IHCs (red) and SGNs (blue) in the three cochlear turns at 58W are shown as percentage. Ratios were calculated as the average of 1-(KO/WT) for each turn. Data are represented as means ± SEM (n=3-7).

**Figure 3.**
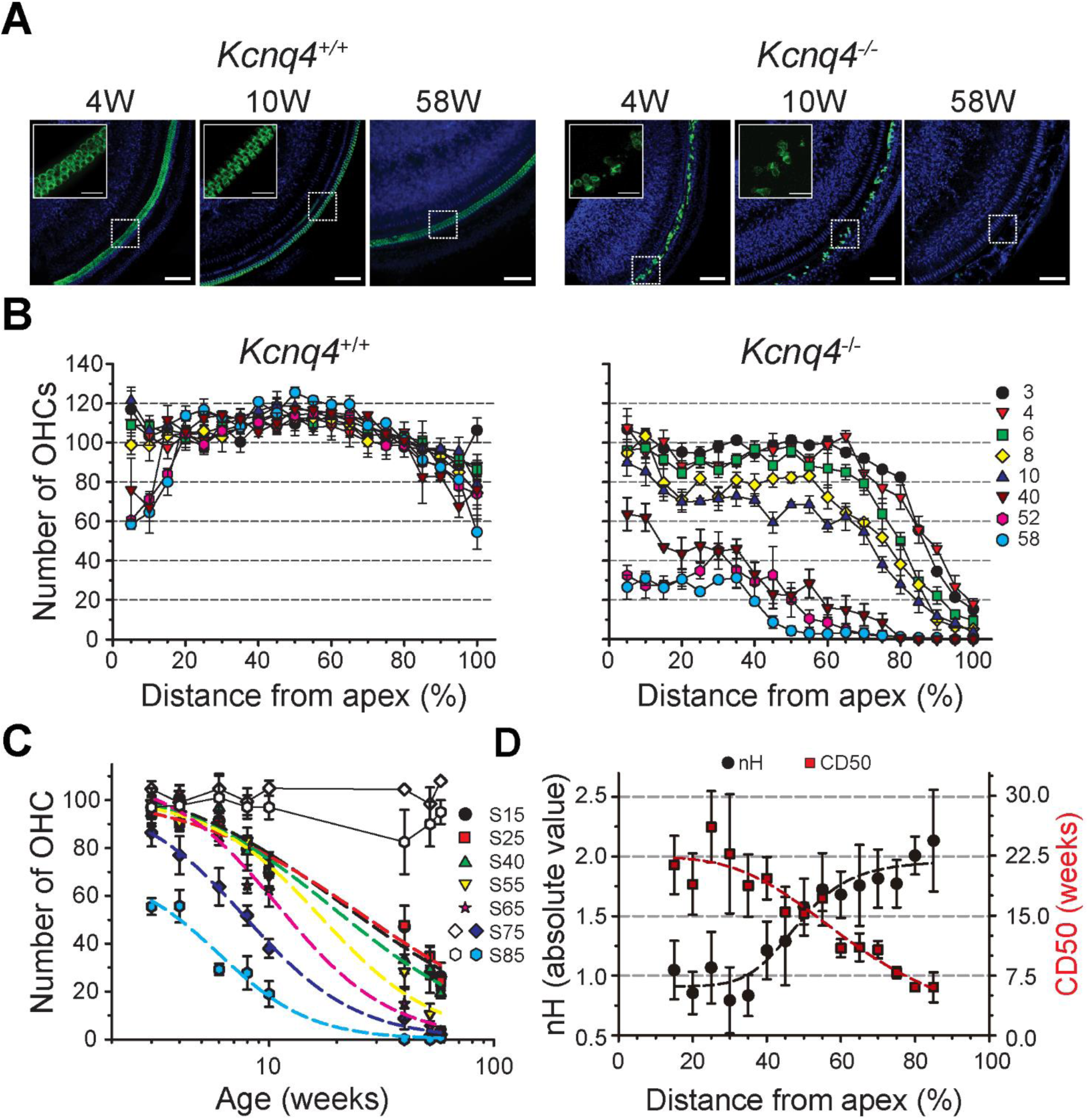
Spatiotemporal analysis of OHC loss *Kcnq4*^-/-^ mice. **(A)** Representative pictures of OHCs from *Kcnq4*^+/+^ (left) and *Kcnq4*^-/-^ (right) mouse cochleae. Whole-mount sections show OHC from basal turns at 4, 10 and 58W labeled with anti-prestin (green) antibody. Scale bar: 75 μm. Nuclei were stained with DAPI (blue). Insets show a higher magnification of the area delimited by the corresponding dashed white lines. Scale bar: 20 μm. Insets corresponding to 58W are shown in Fig. 2A. **(B)** OHC cytocochleograms of *Kcnq4*^+/+^ (left) and *Kcnq4*^-/-^ (right) mice across 3W to 58W range. Values represent the mean number of OHC in each S and age ± SEM (n=3-5). Symbols indicate mouse age (W) and correspond to: 3W: black circle, 4W red inverted triangle, 6W: green square, 8W: yellow diamond, 10W: blue triangle, 40W: brown inverted triangle, 52W pink hexagon, 58W: light blue circle. **(C)** Assessment of OHC loss rates at each segment for *Kcnq4*^-/-^ mice. OHC mean counts were plotted at the indicated S vs. age (taken from panel B). Data were fitted using Hill equation (dashed lines) (see Experimental Procedures). Symbols for the different S are: S15: black circle, S25: red square, S40: green triangle, S55: inverted yellow triangle, S65: pink star, S75: blue diamond and S85: light blue hexagon. Open diamond and hexagon correspond to WT S75 and S85, respectively. Data are represented as means ± SEM (n=3-5) **(D)** Kinetic parameters for the Hill equation corresponding to each segment. Hill coefficients nH (left axis) and CD50 (right axis) were plotted. Symbols correspond to: nH: black circles, CD50: red squares. Data are represented as means ± SEM (n=3-5).

**Figure 4.**
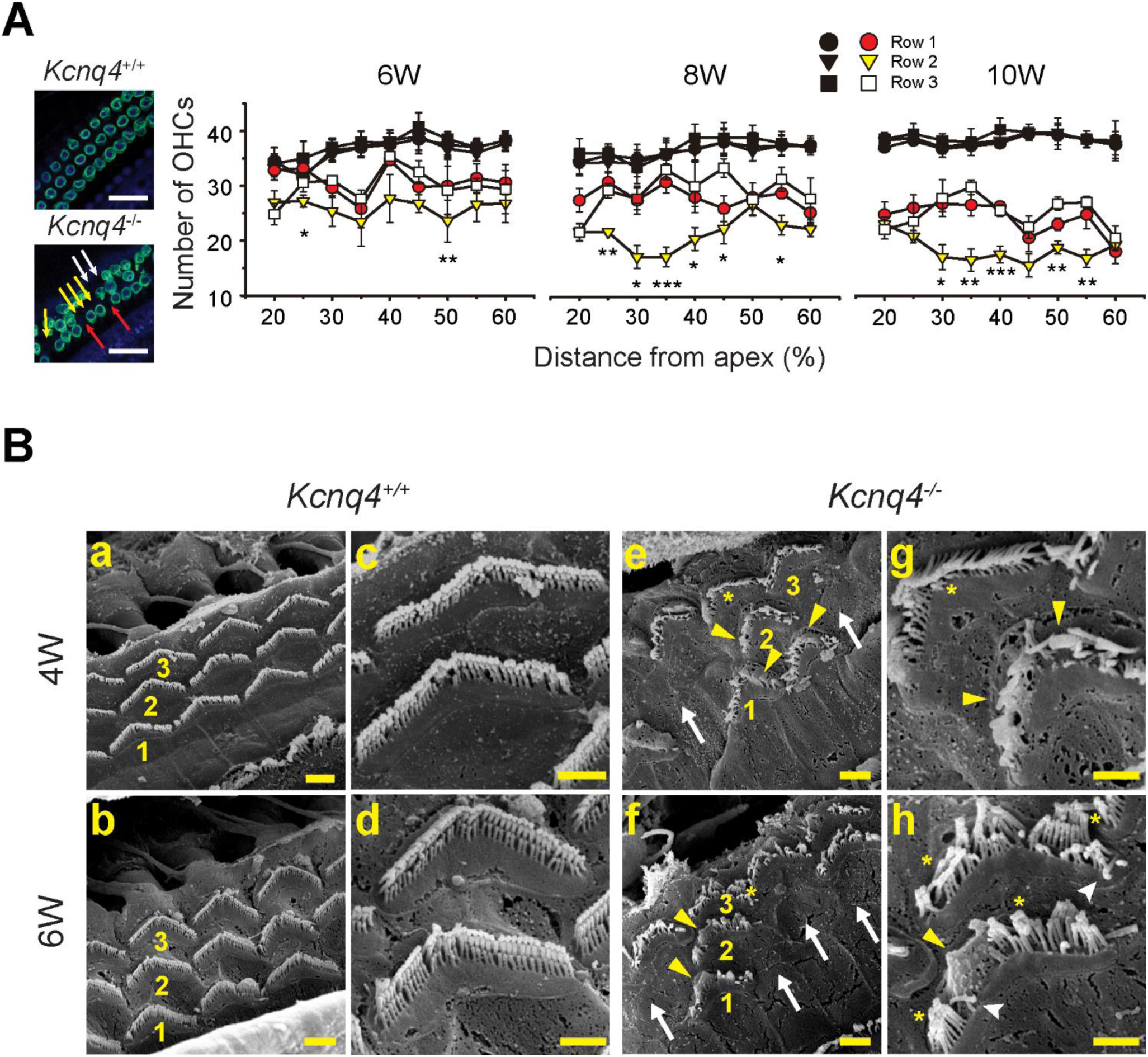
Row architecture and surface ultrastructure of OHCs in young *Kcnq4*^-/-^ mice. **(A)** OHC survival analysis for each of the three rows. The left side shows representative immunofluorescence pictures from 10W *Kcnq4*^+/+^ (upper) and *Kcnq4*^-/-^ mice (lower). OHCs were labeled with anti-prestin (green) and nuclei with DAPI (blue). Arrows point out missing OHCs in row 1 (red), row 2 (yellow) and row 3 (white), in *Kcnq4*^-/-^ mouse. Scale bar, 25 μm. On the right it is displayed average OHC counts for each row in WT (dark symbols) and KO (colored symbols) mice. OHCs were counted separately for row 1 (circle), row 2 (triangle) and row 3 (square) in the S20-S60 range at different ages. Data are represented as means ± SEM from 4-5 independent experiments. For statistical analysis, each row was compared to the other 2, at each segment in both genotypes. No statistical differences were observed among rows for *Kcnq4*^+/+^ animals at all ages. Asterisks indicate statistical differences between row 2 and the other two for *Kcnq4*^-/-^ animals. At 6W; S25, p<0.050 (*) (row1 vs. row2) and S50; p<0.010 (**) (row2 vs. row3). At 8W, S25; p<0.050 (**) (row2 vs. row3); S30, p<0.050 (*) (row2 vs. row3); S35, p<0.001 (***) (row2 vs. row3); S40, p<0.050 (*) (row2 vs. row3); S45, p<0.050 (*) (row2 vs. row3) and S55, p<0.050 (*) (row2 vs. row3). At 10W; S30, p<0.050 (*) (row2 vs. row3); S35, p<0.010 (**) (row2 vs. row3); S40, p<0.001 (***) (row1 vs. row2); S50, p<0.010 (**) (row2 vs. row3) and S55, p<0.010 (**) (row2 vs. row3) (ANOVA). **(B).** Apical surface analysis of hair cells by scanning electron images in WT (a-d) and KO mice (e-h). Pictures at low (a, b, e and f) and high (c, d, g and h) resolution are shown for 4W (upper) and 6W (lower) mice. Internal, middle and external OHC rows are indicated as 1, 2 and 3, respectively. Symbols indicate: white arrows, OHC absence; white arrowheads, floppy stereocilia; yellow asterisks, tip-fused stereocilia and yellow arrowhead indicate whole-fused structures. Scale bar: 2 µm in a, b, e and f and 1 µm in c, d, g and h.

**Figure 5.**
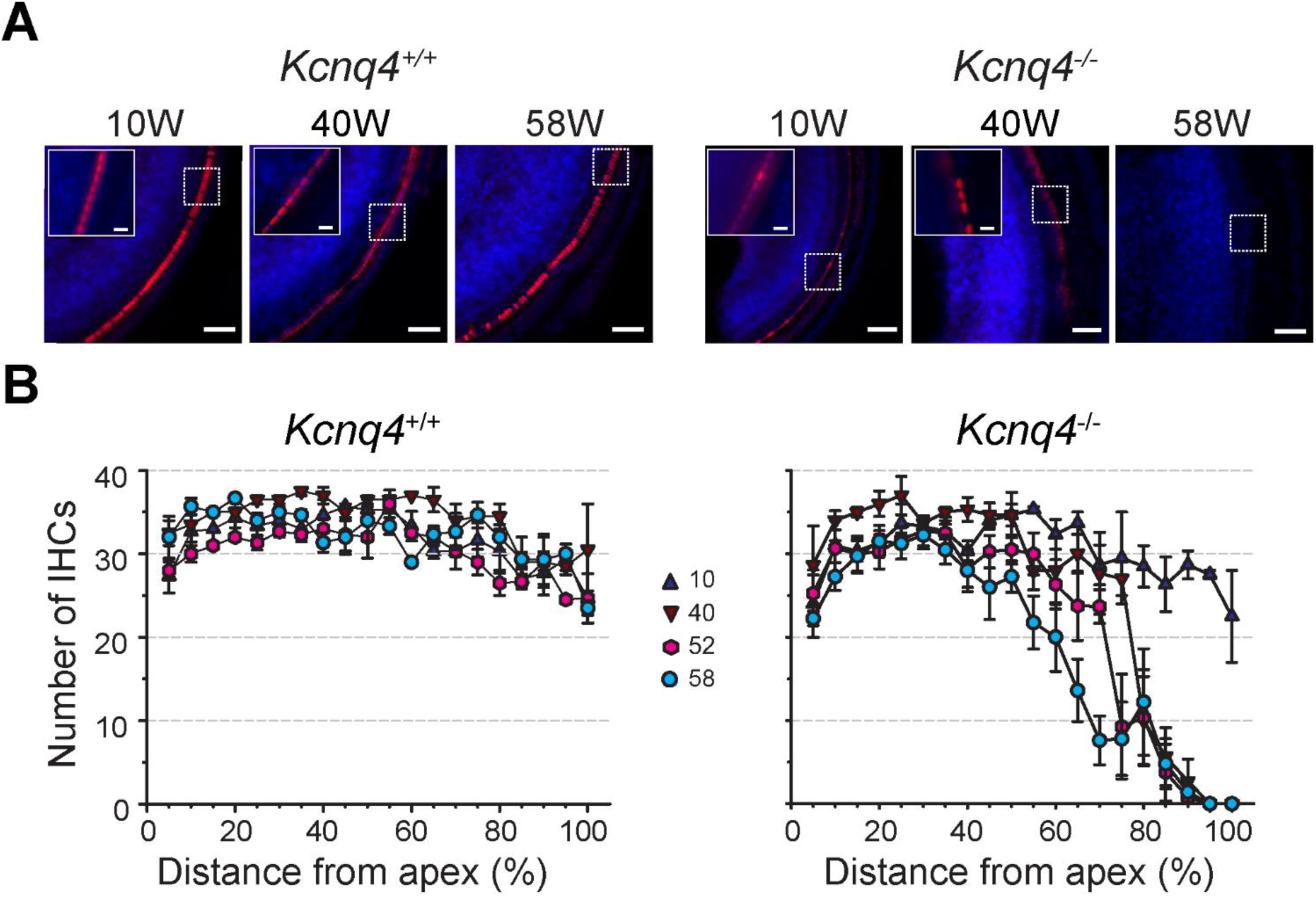
Spatiotemporal analysis of IHC loss in *Kcnq4*^-/-^ mice. **(A)** Representative pictures of WT (left) and KO (right) mice cochleae at 10, 40 and 58W showing IHCs in basal turn. IHCs were labeled using anti-myosin VIIa (red) antibody in cochlear whole-mount preparations. Nuclei were stained with DAPI (blue). Scale bar: 50 μm. Insets show a higher magnification of the area delimited by the corresponding dashed white lines. Inset pictures for 58W mice are shown in Fig. 2A. Scale bar: 10 μm. **(B)** IHC cytocochleograms of *Kcnq4*^+/+^ (left) and *Kcnq4*^-/-^ (right) mice at several ages. Values represent the mean number of IHC in each S and age ± SEM (n=3-5). Symbols indicate mouse age (W) and correspond to 10W: blue triangle, 40W: brown inverted triangle, 52W: pink hexagon, 58W: light blue circle.

**Figure 6.**
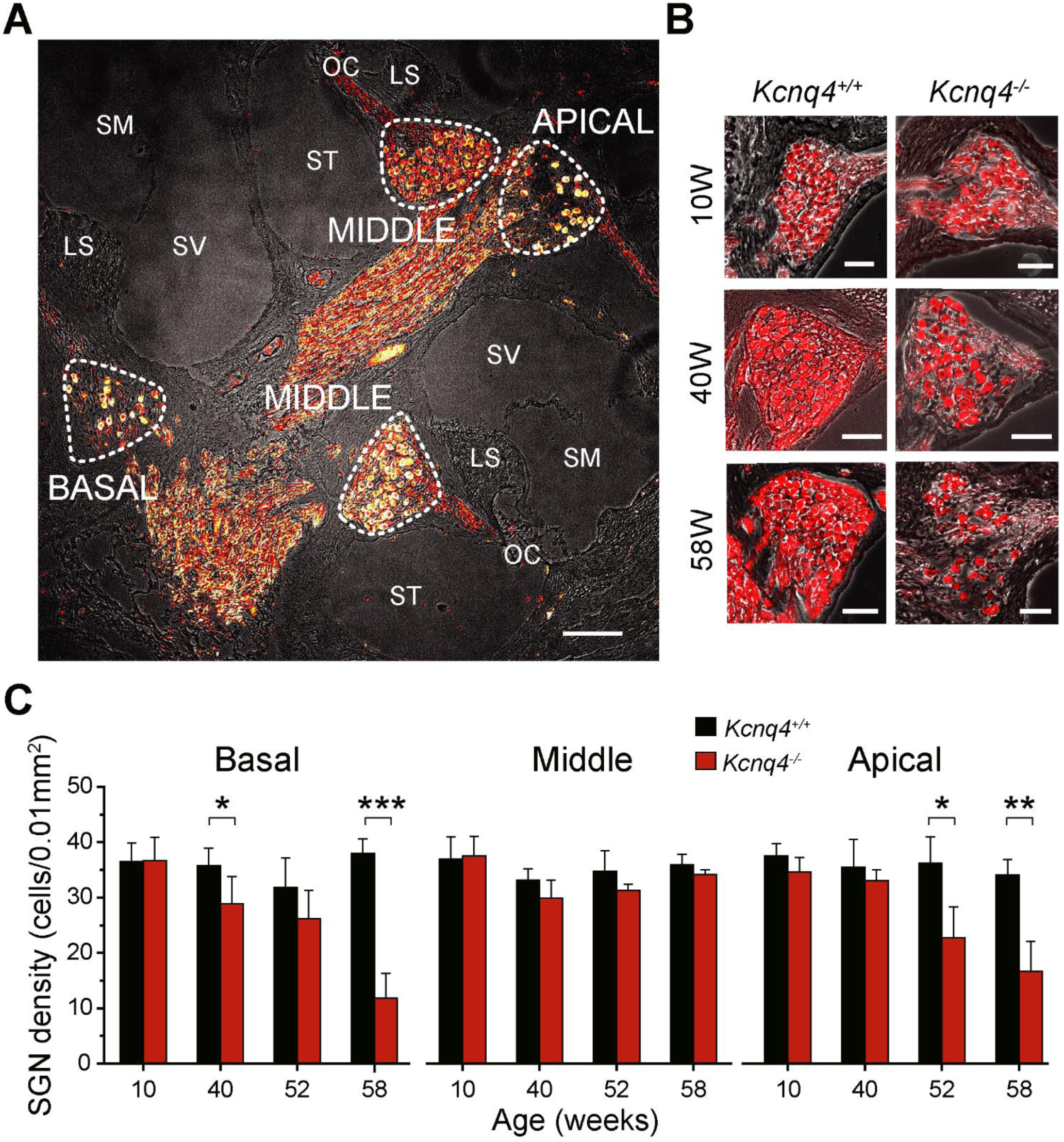
Spiral ganglion neuron loss in *Kcnq4*^-/-^ mice. **(A)** Representative picture of a modiolar cochlear section showing cochlear structures including the three turns of the spiral ganglion. Picture was taken from a 52W *Kcnq4*^-/-^ mice. Neurons were labeled with anti-β-III tubulin (red). White dashed lines delimit spiral ganglia area. β-III tubulin signal was superimposed on the corresponding light transmission phase contrast pictures. Abbreviations are: SV: Scala vestibuli, SM: Scala media, ST: Scala timpani, OC: Organ of Corti, LS: Limbus spiralis. Cochlear turns are also indicated Scale bar: 100 µm. **(B)** Representative pictures showing SGNs in basal turn from *Kcnq4*^+/+^ and *Kcnq4*^-/-^ mice at 10W, 40W and 58W. β-III tubulin-positive neurons are red-colored. Corresponding phase-contrast pictures are also shown. Scale bar: 50 µm. **(C)** SGN density plot for *Kcnq4*^+/+^ and *Kcnq4*^-/-^ mice through age in basal (left), middle (middle) and apical (right) cochlear turns. SGN density values were obtained counting the number of β-III tubulin-positive neurons in the selected area in each cochlear turn at the indicated age. Data are represented as means ± SEM from 3-5 independent experiments. Asterisks indicate statistical differences between *Kcnq4*^+/+^ and *Kcnq4*^-/-^ animals. At basal turn, p<0.050 (*) for 40W and p<0.001 (***) for 58W. At apical turn, p<0.050 (*) for 52W and p<0.010 (**) for 58W; Student´s t test.

**Figure 7.**
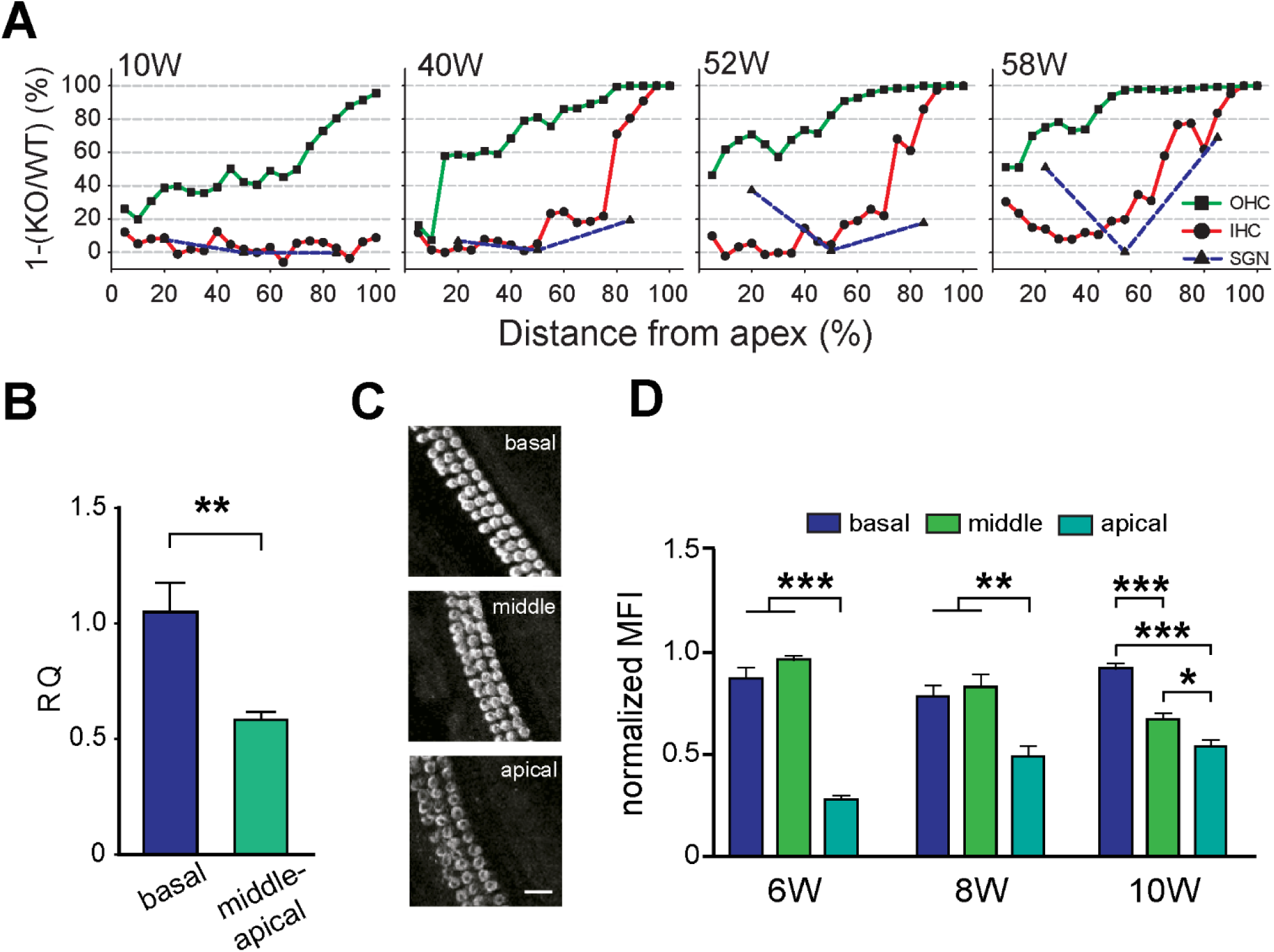
Spatiotemporal correlation between cell loss and KCNQ4 channel expression. **(A)** Increase of OHC (square), IHC (circle) and SGN (triangle) loss from 10W to 58W mice. Cell loss is expressed as 1-survival ratio (KO/WT). Data were calculated from cytocochleograms for OHCs and IHCs and neuronal densities for each cochlear turn for SGNs. **(B)** *Kcnq4* gene expression in WT mouse cochleae. Relative quantification (RQ) of *Kcnq4* mRNA was performed in the basal and middle/apical turns of 3W WT mouse cochleae. Total RNA was extracted and RT-qPCR was performed. The fold change of *Kcnq4* was calculated using 2^-ΔΔCT^ and mRNA expression was referred to *Gapdh* and *Hprt* genes and compared to the basal turn. Data are represented as means ± SEM (n=3-4 pooled animals, 2 independent experiments). P<0.010 (**), Student’s t-test. **(C)** Representative pictures showing KCNQ4 immunolabeled OHCs in the three different cochlear turns at 10W mice. Cochlear whole-mounts were processed separately for basal, middle and apical turns and the fluorescence intensity was quantified in at least 20 cells/turn. Scale bar, 15 μm. **(D)** Mean fluorescence intensity (MFI) quantification of KCNQ4-positive cells in young adult mice. MFI were analyzed with Image J software and values were normalized to their maximum at each age. Data are represented as means ± SEM (n=3). At 6W, ANOVA shows statistical differences between apical to basal or middle turns (p<0.001) (***). At 8W, ANOVA shows statistical differences between apical to basal or middle turns (p<0.010) (**). For 10W, ANOVA shows statistical differences between apical to basal (p<0.001) (***), between apical to middle (p<0.05) (*) and between middle to basal turns (p<0.001) (***).

## RESULTS

### Effect of genetic background on cochlear cell survival

Genetic backgrounds are well known to influence disease phenotype (Doetschman, 2009). Previous studies with *Kcnq4* KO allele were performed on a mixed background between C57BL/6 and 129/SV mouse strains which displayed only OHC disappearance, resembling the initial phase of DFNA2 (Kharkovets et al., 2006). To better reproduce DFNA2 phenotype including early and late phases, we expressed *Kcnq4* KO allele in C3H/HeJ strain. In this last, hearing function has been reported to be well preserved beyond 1 year-old with a low incidence of ARHL (Trune et al., 1996). For this reason, it is a suitable background to evaluate progressive hearing loss in aged mice. Thus, we first analyzed cell survival in 52W WT C3H/HeJ mouse strain to determine normal hair cell loss. Then we contrasted these data with those of age-matched C57BL/6 mouse strain, a known model for ARHL (Davis et al., 2001). We used a standard procedure based on cytocochleographic analyses, which related cell counts with cochlear segments (see Experimental Procedures). OHCs were identified using an anti-prestin antibody and IHCs by anti-Myosin VIIa antibody (Kharkovets et al., 2006, Dallos, 2008, Spitzmaul et al., 2013). Figure 1A (left) depicts a schematic division of the cochlea that was performed for whole-mount of the organ OC. Traversed yellow dashed lines indicate the boundaries of the 5%-length segments (S) in which the entire cochlea was divided and where hair cells were counted (see Experimental Procedures). Both strains showed robust signals for OHCs (green) and IHCs (red) in the three cochlear turns. While C3H/HeJ showed the normal pattern of HC distribution with almost no gaps, C57BL/6 exhibited missing cells in basal and middle turns for both HC types (Fig. 1B). For OHCs, both strains exhibited similar survival patterns in apical and middle cochlear turns (S15 to S60) (Fig. 1C). However, the amount of OHCs in C57BL/6 mice progressively decreased from S65 onwards, with no detectable OHCs in the last four segments. Conversely the C3H/HeJ strain only developed a slight OHC loss at the basal turn. Yet there was a small but significant decrease in OHCs on the first two apical segments compared to C57BL/6 (p<0.05 for S5 and S10, respectively; Student´s t-test) (Fig. 1C). On the other hand, IHC number in C3H/HeJ mice remains relatively constant along the cochlea, while in C57BL/6 mice these cells decrease abruptly in the farthest basal segments (S85-S100) (p<0.05; Student´s t-test) (Fig. 1C, right). Thus, our results indicate a high proportion of HC survival in C3H/HeJ strain, enabling to analyze the role of KCNQ4 channel in this process at least up to one year of age.

### Effect of KCNQ4 deletion on cochlear cell survival in middle-age mice

To have an overview of the impact of KCNQ4 channel absence on cell survival, we determined the number of OHCs, IHCs and SGNs in the three cochlear turns at 58W WT and *Kcnq4*^-/-^ mice (Fig. 2). *Kcnq4*^+/+^ cochlea showed the presence of all 3 rows of OHCs in the three turns with occasional missing cells in apical or basal turns (Fig. 2A), as described in Figure 1. No loss was observed neither for IHCs nor SGNs counts, whose average values were 32.3±3.0 cells/turn for IHCs and 36.0±1.9 cells/0.01 mm^2^ for SGNs (Fig. 2A, left). In contrast, a differential decrease for each cell type in *Kcnq4*^-/-^ mice is observed (Fig. 2A, right). To better understand the spatial progression of cell death, we calculated the cell loss ratio between KO and WT animals in each cochlear turn. We observed a drastic decrease of OHCs, from 60% in apical turn to a complete absence in basal turn (Fig. 2B, green bars). For IHC, we observed a mild cell loss in apical and middle turns, but a severe cell loss in basal turn (∼85%) (Fig. 2B, red bars). For SGNs apical and basal turns were the most affected, with a 50% and a 30% cell survival, respectively, while the middle turn did not exhibit cell loss at all (Fig. 2B, blue bars).

### OHC loss along the Organ of Corti in *Kcnq4*^-/-^ mice

KCNQ4 channel absence impairs K^+^ extrusion from OHCs which would disturb K^+^ recycling in the inner ear, leading to OHC death and tissue degeneration (Kharkovets et al., 2006). We monitored cell survival in cochlear whole-mount preparations from WT and *Kcnq4*^-/-^ mice from 3W to 58W. First we analyzed tissue length to avoid miscalculation of cell counts and found that was similar for both genotypes (p>0.050). Average length values were 5497±190 µm (n=24) and 5465±168 µm (n=26) for *Kcnq4*^*+/+*^ and *Kcnq4*^*-/-*^ mice, respectively.

In order to evaluate the spatial OHC loss exhibited by *Kcnq4*^*-/-*^ mouse, we performed cytocochleographic analyses. The number of OHCs in each 5%-segment were explored and compared between both mouse genotypes at different ages. While in WT animals OHC number remained almost constant, a decrease in OHC counts was observed for *Kcnq4*^-/-^ mice (Fig. 3A, see also Fig. 2A, OHC panel). Cytocochleographic analyses of young WT mice (3W-10W) showed no changes in OHC number throughout the cochlea, except for the basal segments (i.e. >S90), where a slight but still significant decrease in cell number was observed from 4W to 10W (p<0.05) (Fig. 3B, left). In middle-age adult WT mice (40W-58W), OHC survival was similar to that found in young animals within the length range of S20 to S90. These mice showed as well, a significant reduction in the number of OHCs in apical and basal segments (Fig. 3B, left) (p<0.05 for S5, S10, S15 and S100, respectively).

OHC survival was drastically altered in *Kcnq4*^-/-^ mice. At the youngest studied age (3W), OHC counts were slightly lower than in *Kcnq4*^+/+^ until S80, significantly dropping down from this point onwards. Cell number progressively decreased from S85 to S100, exhibiting an 85% OHC loss at the cochlear hook (S95-S100) (Fig. 3B, right). As mice grew old, the number of OHCs in *Kcnq4*^-/-^ mice decreased progressively across the entire cochlear length (Fig. 3B, right). While no significant differences were observed between 3W and 4W, cell survival gradually decreased with age becoming significant at basal segments (S95 at 6W), shifting to apical segments as age increased (Table 1, Fig. 3B). Middle-age adult *Kncq4*^-/-^ mice exhibited a complete absence of OHCs in basal segments from the S80 forward. Middle-to-apical segments appeared to be less sensitive to the loss of the KCNQ4 protein showing a 20-40% of remaining OHCs (Fig. 3B, right).

**TABLE 1.**
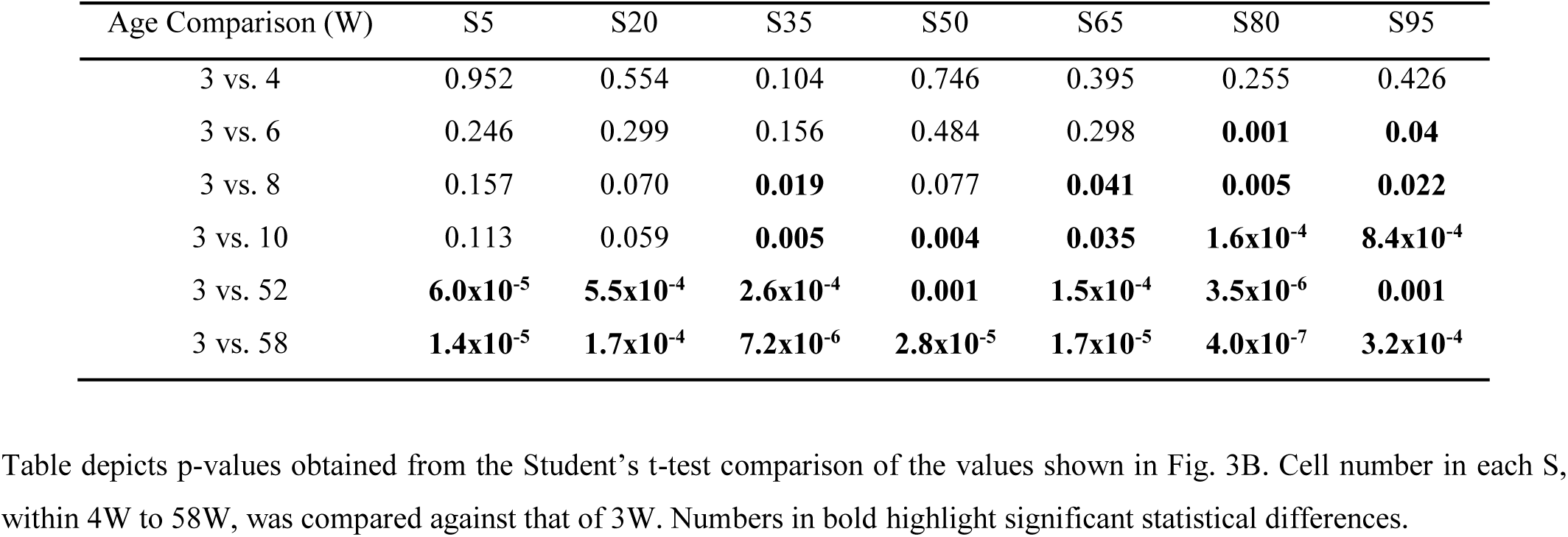
Statistical analyses of OHC number for representative cochlear segments (S) in *Kcnq4*^*-/-*^ mice.

| Age Comparison (W) | S5 | S20 | S35 | S50 | S65 | S80 | S95 |
| --- | --- | --- | --- | --- | --- | --- | --- |
| 3 vs. 4 | 0.952 | 0.554 | 0.104 | 0.746 | 0.395 | 0.255 | 0.426 |
| 3 vs. 6 | 0.246 | 0.299 | 0.156 | 0.484 | 0.298 | **0.001** | **0.04** |
| 3 vs. 8 | 0.157 | 0.070 | **0.019** | 0.077 | **0.041** | **0.005** | **0.022** |
| 3 vs. 10 | 0.113 | 0.059 | **0.005** | **0.004** | **0.035** | **1.6x10<sup>-4</sup>** | **8.4x10<sup>-4</sup>** |
| 3 vs. 52 | **6.0x10<sup>-5</sup>** | **5.5x10<sup>-4</sup>** | **2.6x10<sup>-4</sup>** | **0.001** | **1.5x10<sup>-4</sup>** | **3.5x10<sup>-6</sup>** | **0.001** |
| 3 vs. 58 | **1.4x10<sup>-5</sup>** | **1.7x10<sup>-4</sup>** | **7.2x10<sup>-6</sup>** | **2.8x10<sup>-5</sup>** | **1.7x10<sup>-5</sup>** | **4.0x10<sup>-7</sup>** | **3.2x10<sup>-4</sup>** |

In order to evaluate the kinetic of OHC loss in cochlear turns, we analyzed the rate of cell loss at every S by plotting the number of OHCs against mouse age (Fig. 3C). While in *Kcnq4*^+/+^ mice the number of OHCs remained constant over time (Fig. 3C, open symbols), it decayed in *Kcnq4*^-/-^ animals (Fig. 3C, closed symbols). The loss of OHC was slow at the youngest ages but increased drastically in the following weeks until cells degenerated completely older ages. Most apical segments (S5 and S10, not shown) did not follow this pattern, probably due to an overestimation of cell death resulting from the slight age-dependent HL observed also in *Kcnq4*^+/+^ mice (see Fig. 3B). To describe the kinetic of cell loss we used the Hill equation to fit our temporal data for each S. Slopes for apical to middle segments (S15 to S40) were less pronounced than those observed in basal segments (Fig. 3C). In order to visualize the behavior of OHC loss rates, we plotted Hill coefficients and the CD50s obtained from fitted curves for *Kcnq4*^-/-^ mice at each cochlear S (Fig. 3D). Hill coefficient progressively increased from apical (nH ∼0.9) to basal (nH ∼2.0) segments in a sigmoidal shape. Conversely, we observed that CD50 decreased with increasing distance from the apex, from an initial value of around 22 weeks (S15) to a value of 7 weeks at S85 (Fig. 3D). Changes in coefficient values indicate that kinetic of OHC loss is not homogenous along cochlear segments, speeding up from apical to basal turns. For the same age, OHC number will decrease in apical segments proportionally to 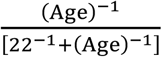 while in basal segments it will be 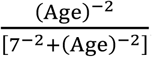.

### Alteration of OHC row architecture and stereociliar structure in *Kcnq4*^-/-^ mice

To get insights into OHC degeneration, we analyzed the time-course pattern of OHC survival in each individual row. Cochleographic studies were carried out in the S20-S60 range for young animals (6W to 10W) since OHC degeneration from these cochlear regions is ongoing within this timeframe, but cell row architecture is still well preserved (Fig. 4A, left). In *Kcnq4*^+/+^ mice, we did not observe differences among OHC counts for each row, exhibiting an average value of ∼37 cells/row for each segment (Fig. 4A). In agreement with the results shown above, the OHC number for each row decreased with age in *Kcnq4*^-/-^ animals. However, OHC loss was not equivalent among the three rows (Fig. 4A). Indeed, the highest decrease was observed for the central row 2. At 6W, only two segments of row 2 showed statistically significant differences respect to the internal row 1 and the external row 3: S25 and S50 (p<0.05). The rest of the segments in row 2 exhibited lower cell counts than row 1 and 3, which did not reach statistical significance, suggesting a trend towards higher cell death rate (Fig. 4A). At 8W and 10W, differences between row 2 and the others became more evident. Indeed, at 8W and 10W, 5 to 6 segments out of 9 in row 2 showed significant differences (Fig. 4A).

Due to the differential cell loss among rows and to better understand the process of tissue degeneration, we studied apical surface by SEM (Fig. 4B). As expected, WT mice inspection at 4W and 6W showed the characteristic 3-row distribution of OHCs without spaces between them (Fig. 4Ba and 4Bb). In addition, individual cells bared the typically polarized hair bundle structure, where the upper surface ends up with ordered and neat rows of stereocilia, arranged in a 3-step staircase profile, exhibiting the characteristic W shape (Fig. 4Bc and 4Bd). On the other hand, basal cochlear segments of *Kcnq4*^*-/-*^ mice, even at 4W, showed clear alterations in tissue arrangement. The typical 3-row pattern was still recognizable in some regions but not clearly defined in others (Fig. 4Be and 4Bf). Cell loss became apparent at this age because several hair bundles were missing and replaced by scars on tissue surface. OHC loss was even greater at 6W (see white arrows in Fig. 4Be and 4Bf), being these results in agreement with those described above (see Fig. 3 and 4A).

In addition to cell loss, hair bundle structure in the still surviving cells exhibited evident damage showing a disorganized pattern (Fig. 4Bg and 4Bh). Figure 4B allowed to identify several steps of the degenerative process, starting from the tip-fusion of the 3 stereociliar rows (asterisks), progressing to “walls” of fused stereocilia (yellow arrowhead) in which individual ones are no longer recognizable. Occasionally, it was also possible to observe floppy stereocilia (white arrowhead). Fusion processes can span from small to big portions of the bundle. In some cases, stereociliar fusion between two different bundles could be observed. So, OHCs not only displayed cell loss but also several alterations in their apical structure. Stereociliar alterations were not observed for IHCs in this time period (not shown).

### Degeneration of IHCs also occurs in KCNQ4 absence

As we did for the OHC degeneration study, we used cytocochleographic analyses to evaluate IHC loss from WT and KO mice. For this we evaluated IHC survival in young and middle-age adult animals. *Kcnq4*^*+/+*^ mice exhibited a continuous single row of IHCs in basal turn at all tested ages (Fig. 5A, left). On the contrary, in *Kcnq4*^-/-^ mice, loss of IHCs is evident only in adult animals (Fig. 5A, right). *Kcnq4*^*+/+*^ mice showed a mean number of ∼33 IHC/S, remaining constant across the S10 to S80 range for all ages. However, in basal segments (>S80), the number of IHCs was statistically lower at all ages (p<0.05) (Fig. 5B, left). On the contrary, in *Kcnq4*^-*/-*^ mice each age showed different IHC counts along the cochlea. Indeed, while in younger mice (10W) the number of IHC remained practically constant and similar to that of WT, middle-age adult mice showed a strong decrease of IHC number. 40W animals displayed differences that reach significance from S80 onwards (p<0.05), showing an initial 60% decrease in cell number, respect to the apical segments (Fig. 5B, right). From this age forward, IHC loss exhibited a time-dependent increase in cell death progressing from basal to middle turns (Fig. 5B, right). Moreover, at S75, IHC counts became lower than 10 cells/S reaching a complete IHC loss in basal segments in 40 to 58W animals (p<0.05). 58W KO animals evidenced cell loss in apical segments that was not observed in *Kcnq4*^+/+^ mice (Fig. 5B).

We also performed kinetic studies for IHCs. Contrary to OHCs, IHC loss is only observed from S55 onward, so we studied this range. Cell loss developed abruptly with age for each segment, exhibiting high nH values, such as 4.9, 11 and 24 for S60, S65 and S70, respectively, when fitted with the Hill equation. Due to the low number of living cells in the most basal segments, we were unable to fit representative curves for S80 to S100. The kinetic of cell loss for the fitted segments developed very fast with age, progressing exclusively from basal to middle turns up to 58W.

### Spiral ganglion neurons also degenerate in *Kcnq4*^-/-^ mice

We also analyzed SGN survival in mice lacking KCNQ4 channel expression on modiolar cochlear sections. Neuronal density was determined in spiral ganglia from basal, middle and apical turns at different ages (Fig. 6A, dashed lines). WT mice displayed a high number of neurons in each ganglion turn at all ages while *Kcnq4*^-/-^ animals exhibited a decrease in neuron number in basal and apical ganglion turns at 58W (Fig. 6B). Estimation of neuronal densities showed that, *Kcnq4*^+/+^ mice exhibited an average value of 36-39 neurons/0.01mm^2^ for the three cochlear segments at 10W. This value remained practically constant through age for the different segments (Fig. 6C). On the contrary, in *Kcnq4*^-/-^ animals, we determined a neuronal loss that varies differentially with ganglion localization. A clear neuronal loss was seen in basal and apical turns in KO mice older than 1 year (Fig. 6C). Neuron density gradually decreased for basal and apical turns as mice grew old, while it remained almost invariable for middle segments. Although a great variability was obtained for neuronal density estimations due to unevenness in cochlear slices, a clear tendency to a decrease in neuron counts was observed at 40W in the basal segment that increased with age, reaching a 60% neuronal loss at 58W mice (p<0.05) (Fig. 6C). Apical neurons from *Kcnq4*^-/-^ mice also degenerate although differences became significant only at 1 year-old (p<0.05). For middle segments, neurons were well preserved showing similar counts to WT mice up to 58W (Fig. 6C). In summary, in *Kcnq4*^-/-^ mice, SGNs also degenerate but later than OHCs and timely-tuned to IHC loss.

### Correlation between cell loss time-course and expression of KCNQ4 channel

To expose the differences in the time-course of IHC, OHC and SGN loss, we calculated the ratio between *Kcnq4*^-/-^ and WT animals for each cell type at four different ages (Fig. 7A). We plotted cytocochleogram loss ratios for IHCs and OHCs, while for SGNs the representative values obtained for apical, basal and middle turns were plotted at S20, S50 and S85, respectively. While OHC loss is observed at all periods, increasing with age, IHCs started at 40W only after S55 (Fig. 7A). SGN disappearance slightly started at basal turn in 40W mice, increasing both basal and apical turn in older mice. A clear difference in the spatial and temporal pattern of cell loss was evident between IHCs and OHCs. For the same age, a similar cell loss proportion (e.g. 40%) is reached at S75-80 for IHCs and S10-15 for OHCs, indicating that, when IHC loss is present at basal segment, OHC degeneration had already reached middle to apical segments (Fig. 7A). Besides, we observed a shift in the time-course. For example, comparing the same segment (e.g. S80), IHCs reached an 80% cell loss at 40W while the same proportion of OHCs were obtained at 10W (Fig. 7A). Thus, there was a delay of IHC degeneration of about 30 weeks compared to OHC degeneration. SGN degeneration followed a similar time-course pattern than IHCs (Fig. 7A).

Then we analyzed KCNQ4 channel expression in C3H/HeJ WT mice. Using reverse transcription followed by qPCR, we evaluated the expression level of *Kcnq4* mRNA in basal and middle/apical cochlear turns of WT young mice. As shown in Fig. 7B, middle/apical turns exhibited around 50% less amounts of *Kcnq4* mRNA compared to the basal turn (p<0.010).

The expression of the KCNQ4 protein was evaluated by MFI level measurements performed on cochlear whole-mounts. KCNQ4 signal was clearly detected in the three OHC rows while no signal was observed neither for IHCs nor for SGNs (Fig. 7C). Fluorescent signals from the basal, middle and apical turns were compared for young mice. Similar fluorescence intensities were detected in the three rows of OHCs for the same turn, however, it varied with the different turns (Fig. 7C). Fluorescent signal was robust in basal and middle turns, but it was weaker in cells of the apical turn. To quantify the KCNQ4 expression level, we measured MFI levels in young animals. As shown in Figure 7D, the apical turn exhibited a 40 to 60% decrease of the normalized MFI at all ages, when compared to basal turn (p<0.05 for 6W, 8W and 10W, respectively). Besides, at 10W, a significant decrease of fluorescence (about 20%) was observed for middle turn compared to the basal one (p<0.05) and this KCNQ4 channel expression gradient remains similar as mice grew old (not shown).

## DISCUSSION

We investigated the cellular basis of DFNA2 hearing loss by analyzing the cochlear cell survival using the C3H/HeJ mouse strain which carries the *Kcnq4* KO allele. Expression of this allele in a mixed background did not resemble the ultimate phase of DFNA2 (Kharkovets et al., 2006). However, using this inbred mouse strain, we were able to gather many more features better resembling the cellular alterations displayed by DFNA2 patients. Genetic background is very important for phenotype expressivity which can be modulated by modifier genes, increasing or decreasing it (Doetschman, 2009). Inbred strains have a better control of these genes but they must be selected carefully. C57BL/6, a very common inbred strain, is a model for ARHL which develops IHC and OHC loss with age (Kane et al., 2012). The C3H/HeJ mouse background offers advantages for our studies of DFNA2 expressivity over other strains because cell survival is very high, even at advanced ages, keeping age-related hair cell death to a minimum (Trune et al., 1996).

### Time pattern of OHC, IHC and SGN degeneration in the DFNA2-like mouse model

Previous analysis of KCNQ4 KO mouse phenotype showed misfunction and degeneration only of OHCs, related to the first phase of DFNA2 hearing loss (Kharkovets et al., 2006). In our current study, besides OHC degeneration, we also stablished the loss of IHCs and SGNs in mouse lacking KCNQ4 channel expression. However, these cells die much later than OHCs. In our experiments, we detected a significant IHC death by week 40 that was restricted to the basal turn. The proportion of cell loss is very high in basal regions with a complete absence of IHCs at the hook after 40W. Additionally, 1 year-old mice had a slight but significant decrease in cell survival in the most apical segments (around 30%). Thus, this data indicates that middle-age adult KO mice in the C3H/HeJ strain, will suffer from a deep HL to high-frequency sounds and could be linked to profound deafness observed in aged DFNA2 patients (De Leenheer et al., 2002, Dominguez and Dodson, 2012). We also found loss of SGNs, starting in middle-age adults, mostly in parallel with IHC loss. It progresses with age in basal turns, but also in the most apical segments. Distinctly, neuron degeneration was not present in middle turns up to 58W mice. Although the analysis for SGNs was not so detailed due to technical limitations, we detected a strong reduction in the number of neurons. Neuronal loss in apical and basal turns mostly parallels that of IHC, suggesting a link between them. Although cell degeneration is caused by the absence of KCNQ4 channel, cellular death cannot be totally attributed to its lacking.

### DFNA2 hearing loss and KCNQ4 channel expression

Cochlear expression of KCNQ4 protein in C3H/HeJ WT mice was detected only in OHCs with a decreasing gradient of channel expression from basal to apical turns in young and adult mice, in accordance with others (Mammano and Ashmore, 1996, Ruttiger et al., 2004, Kharkovets et al., 2006, Winter et al., 2006, Mustapha et al., 2009, Jaumann et al., 2012, Takahashi et al., 2018). The expression gradient is correlated with the progression of OHC death obtained in our experiments. This result can be associated with the initial reduction of hearing sensitivity (<60 dB) at high frequencies, that progress to middle and low frequencies with age in DFNA2 patients (Dominguez and Dodson, 2012). Also, ototoxic drugs that alter KCNQ4 channel function exhibit a similar degenerative profile, where OHCs from basal turns are more sensitive to their deleterious effects (Fausti et al., 1984, Leitner et al., 2011, Sheppard et al., 2015). However, this expression pattern differs from that reported by others. Beisel et al (Beisel et al., 2005) found that KCNQ4 is highly expressed in OHCs from the apical turn, decreasing towards the basal hook. Besides, they detected expression of KCNQ4 in IHCs and SGNs, which was higher in basal than apical turns. By contrast, our IF experiments could not detect KCNQ4 expression neither in IHCs nor SGNs. The reason for these differences is not clear and they could be generated by the genetic backgrounds employed in each case. Although we could not detect expression of KCNQ4 in IHCs, several other experimental approaches suggest its presence in this cell type (Marcotti et al., 2003, Oliver et al., 2003, Kharkovets et al., 2006). Channel expression seems to be restricted to the neck of the IHC, along with the BK channel (Oliver et al., 2003). KCNQ4 is responsible for the I_K,n_ current of IHCs contributing to restore membrane potential since its absence generates a slight depolarization (Marcotti et al., 2003, Oliver et al., 2003, Beisel et al., 2005, Kharkovets et al., 2006).

Our results indicate that KCNQ4 plays differential roles in cell survival: for OHCs it contributes to short-to middle-term while for IHCs to long-term survival. In OHCs it is the main potassium extruder while in IHCs it participates in restoring membrane potential (Oliver et al., 2003, Kharkovets et al., 2006). In the first case, its absence generates a 15 mV depolarization. On the other hand, KCNQ4 absence in IHCs slightly depolarizes them (3-5 mV), generating a long-term cumulative deleterious effect, leading to cell death 30 weeks later than OHCs in our model.

The expression of KCNQ4 channel in SGNs is contradictory. Although we did not find it by immunoflurescence, its expression had been reported by others (Beisel et al., 2005). These neurons exhibit an M-current generated by KCNQ channels (Lv et al., 2010). KCNQ2 and KCNQ3 subunits were reported in SGNs (Jin et al., 2009). Then, the molecular composition of the M-current detected in these studies cannot be ascribed to KCNQ4 channel and we cannot assert the possibility of cell death due to the lack of this channel. It is still unknown whether neuronal degeneration in KO mouse occurs as a primary event due to KCNQ4 absence or if it is a consequence of IHC loss. There is evidence indicating that SGN survival depends on trophic support provided by IHCs (Gillespie and Shepherd, 2005) and supporting cells (Stankovic et al., 2004, Sugawara et al., 2005, Sugawara et al., 2007). The temporal correlation observed between IHC and SGN death links their degenerative process, at least in apical and basal turns.

As synaptic disconnection of SGNs has been postulated as an earlier event triggering neuron degeneration (Liberman, 2017), the loss of IHCs observed in adult mice would impact on neuron survival. In summary, SGN death is a later event in mouse lacking KNCQ4 expression and its cause cannot be certainly attributed to this channel function, but probably to IHC disappearance.

### HC death kinetics and neuronal loss significance to DFNA2

The loss of OHCs has been reported previously, but the process is not fully understood (Kharkovets et al., 2006, Dominguez and Dodson, 2012). For this reason, we thoroughly track the cell death kinetics for both hair cell types. We constructed cytocochleograms for OHC and IHC survival at different ages. This tool enables not only cell count normalization throughout all studied ages, but it also allows to compare HL among species (e.g. humans) (Viberg and Canlon, 2004). For OHCs, our spatiotemporal analysis revealed that all segments develop cell degeneration, although at different rates. Surprisingly by week 3, basal segments already showed a pronounced cell loss. As this was the youngest age tested, we assume OHC degeneration had already begun, considering that KCNQ4 channel is not required for cell differentiation or maturation (Bulankina and Moser, 2012). Cell survival was tracked during several weeks in young and middle-age animals allowing us to determine cell death rates for each segment. We observed that the basal turn had the highest degeneration rate while for the apical turn was the opposite. When we plotted cell survival for each segment as a function of age, we observed that cell loss was not linear in any cochlear segment. Considering that OHCs from KO and KI mouse models for KCNQ4 are chronically depolarized (Kharkovets et al., 2006), our results suggest that at the initial steps of hearing loss, OHCs are able to endure the altered resting membrane potential for some weeks before starting to die abruptly. Interestingly, the rates for cell death are not uniform. To obtain rate trends we fitted our data with the Hill equation. Kinetic parameters indicated that cell loss in the basal turns occurs faster than in the apical turns. Our data is in agreement with the differential rates observed in DFNA2 patients where hearing loss is faster for high-than for low-frequency sounds (Dominguez and Dodson, 2012). Although the progression of OHC loss in KO mouse is much faster than that observed in heterozygous KI mouse (Kharkovets et al., 2006), our results support this pattern of tissue degeneration. In consequence, our data indicate that the expression level of KCNQ4 is relevant to the potassium extrusion function in OHCs because there is a direct correlation between channel expression levels and cell death kinetics.

On the other hand, in middle-age mouse, IHC loss follows a different pattern compared to OHCs exhibiting a drastic reduction in cell number. For example, it takes fewer segments to obtain a high percentage of cell loss for IHC than for OHCs. For this reason, we could only analyze the kinetic of cell loss for segments greater than S55. The loss of IHC was not previously reported in others *Kcnq4* transgenic models. However, IHCs have been suggested to be involved in the process of profound hearing loss reported in aged DFNA2 patients (Dominguez and Dodson, 2012). Our findings show, for the first time, that these cells participate in the progression of deafness since its absence increase hearing threshold more than 70 dB (Willott, 2001). According to our results, this process starts at basal segments, where high-frequency sounds are detected, progressing to middle segments with age, similar to that observed for DFNA2. Surprisingly, for older animals (>1 year-old), we also determined IHC degeneration in apical regions that would impair hearing sensitivity for low-frequency sounds. The heterozygous KI mouse retains 1/16 of the I_K,n_ current, a condition that should occur in DFNA2 patients. The first phase of the disease starts at ∼15 years old, while the second phase appears after 65 years old. The degenerative process is much faster in KO than in KI mice, advancing OHC disappearance (Kharkovets et al., 2006). Therefore, as our model lacks the I_K,n_ current completely, a shift in the life period where HC degeneration occurs, is expected.

Surprisingly we found degeneration of SGNs in basal and apical turns of mice older than 1 year. The participation of SGNs in DFNA2 progression has not been reported. Probably, SGN death is secondary to IHC loss, but its death suggests a progression of neuron degeneration to the CNS, that could affect sound interpretation.

### Tisular and structural alterations of OHCs

We determined a differential OHC row degeneration with age. Although most OHCs die, the middle row showed a significantly higher sensitivity to KCNQ4 absence. The contribution of individual OHC rows to hearing has not been clearly established yet. Modeling predictions indicate that hearing amplification requires proper integrity of all three rows of OHCs. However, a controlled exposure to styrene which produces a specific loss of the outer row had no significant effect on cochlear sensitivity (Chen et al., 2008, Murakoshi et al., 2015). Although we have not measured the functional properties of OHCs, we observed alterations in hair bundle structures at early stages that must affect amplificatory function and contribute to hearing impairment before cell death and replacement by supporting cells. Stereociliar alterations are an early event observed during processes that generate cochlear damage (Slepecky, 1986, Harrison, 2012). Chronic depolarization of OHCs generated by I_K,n_ absence would keep mechanical work exerted by prestin constantly active leading to cell overstimulation and damage in a similar fashion to NIHL (Kao et al., 2013). Besides, *Kcnq4* gene polymorphisms have been associated with ARHL suggesting that a decrease in KCNQ4 channel function may contribute to its subtle and slow progressive hearing impairment (Van Eyken et al., 2006). Then, KCNQ4 KO mouse would be a suitable model to investigate molecular mechanisms and to test drugs to help or prevent several HL pathologies. Sustained depolarization of cell membrane would pose continuous cellular stress that, depending on its intensity will gate cell death sooner or later. In consequence, a better comprehension on the molecular mechanisms involved in KCNQ4 channel function would impact in our understanding of these diseases and their treatments.

## CONCLUSIONS

Our results provide evidence to explain at cellular level the progression of human DFNA2 from its initial steps to the profound deafness observed in aged patients (older than 70 year-old). The early stage of HL is compatible with progressive OHC death from basal to apical turns and the later stage correlates with IHC and SGN disappearance. Our data showed that all cells involved in the initial steps of sound transduction are differentially affected by KCNQ4 absence, posing the intriguing possibility of a neuronal contribution on the progression of hearing loss.

## ACKNOWLEDGMENTS

Our special thanks to Prof. Thomas Jentsch (Max-Delbrück-Centrum für Molekulare Medizin and Leibniz-Institut für Molekulare Pharmakologie, Berlin, Germany) for generously providing *Kcnq4*^-/-^ animals. This work was supported by grants from Agencia Nacional de Promoción Científica y Tecnológica (PICT 2016 N°0260) and Universidad Nacional del Sur (PGI N°24/B262) to GS.

## Abbreviations

OC: organ of Corti
CNS: central nervous system
HCs: hair cells
OHC: outer hair cell
IHC: inner hair cell
SGN: spiral ganglion neuron
WT: wild-type
KO: knock-out
KI: knock-in
*Kcnq4*^*+/+*^: KCNQ4 WT mice
*Kcnq4*^*-/-*^: KCNQ4 KO mice
HL: hearing loss
W: postnatal weeks-old
ARHL: age-related hearing loss
NIHL: noise-induced hearing loss
MFI: mean fluorescence intensity
SEM: scanning electron microscopy
S: 5%-length segment
K^+^: potassium ion
ABR: auditory brainstem response.

